# Improving multi-site autism classification based on site-dependence minimisation and second-order functional connectivity

**DOI:** 10.1101/2020.02.01.930073

**Authors:** Mwiza Kunda, Shuo Zhou, Gaolang Gong, Haiping Lu

**Author notes:** Corresponding author (Haiping Lu). Email addresses (Mwiza Kunda), (Shuo Zhou), (Gaolang Gong).

## Abstract

Autism spectrum disorder (ASD) has no objective diagnosis method despite having a high prevalence. Machine learning has been widely used to develop classification models for ASD using neuroimaging data. Recently, studies have shifted towards using large multi-site neuroimaging datasets to boost the clinical applicability and statistical power of results. However, the classification performance is hindered by the heterogeneous nature of agglomerative datasets. In this paper, we propose new methods for multi-site autism classification using the Autism Brain Imaging Data Exchange (ABIDE) dataset. We firstly propose a new second-order measure of functional connectivity (FC) named as Tangent Pearson embedding to extract better features for classification. Then we assess the statistical dependence between acquisition sites and FC features, and apply a domain adaptation approach to minimise the site dependence of FC features to improve classification. Our analysis shows that 1) statistical dependence between site and FC features is statistically significant at the 5% level, and 2) extracting second-order features from neuroimaging data and minimising their site dependence can improve over state-of-the-art classification results on the ABIDE dataset, achieving a classification accuracy of 73%.

## 1. Introduction

Autism spectrum disorder (ASD) refers to a lifelong neurodevelopmental disorder characterised by a wide range of symptoms, skills and levels of disability, such as deficits in social communication, interaction and the presentation of repetitive patterns of behaviour or restricted interests (Baio, 2014). Until now, there is no known objective method for autism diagnosis with progress mainly challenged by the significant behavioural heterogeneity and wide array of neuroanatomical abnormalities that can be exhibited between patients with autism (Zielinski et al., 2014; Zwaigenbaum & Penner, 2018).

Non-invasive brain imaging techniques such as magnetic resonance imaging (MRI) have been used to discover structural or functional differences between ASD and typical control (TC) subjects. In particular, resting-state functional MRI (rs-fMRI) has achieved promising results when utilised with machine learning (ML) models for classifying ASD and TC subjects (Du et al., 2018). However, the clinical generalisability of most studies using rs-fMRI data for autism classification is debatable since the sample sizes used are small, unlikely to cover a wide spectrum of autism and its heterogeneity. These small sample sizes are due to the time and cost constraint imposed upon *single-site* studies acquiring rs-fMRI using a single fMRI scanner and subject acquisition protocol.

To improve the statistical power and generalisability of neuroimaging studies, the Autism Brain Imaging Data Exchange (ABIDE) initiative has aggregated data from multiple sites across the world, creating datasets much larger than those used in single-site studies (Di Martino et al., 2014). The ABIDE dataset is composed of rs-fMRI and phenotypic data from 20 different international sites, leading to a heterogeneous sample with size of over 1000 ASD and TC subjects. While it presents a great potential for the extraction of functional biomarkers for autism classification, its multi-site and multi-protocol aspects bring along significant patient heterogeneity, statistical noise and experimental differences in the rs-fMRI data, making the classification task much more challenging (Abraham et al. (2017). Recent works have employed different ML methods, such as recurrent neural networks (RNN), graph convolutional neural networks (GCN) and denoising autoencoders (Dvornek et al., 2018; Heinsfeld et al., 2018; Ktena et al., 2018; Parisot et al., 2018). However, despite the complexity in patterns that these methods can generally capture, the difference in their top classification results on ABIDE fall less than 1%, with the highest achieved accuracy being 70.4% by the GCN model developed by Parisot et al. (2018).

This paper investigates two research questions that can potentially improve multi-site autism classification.

- **Between-site heterogeneity:** *how can we effectively account for the experimental differences in the ABIDE rs-fMRI data?* Previous studies on ABIDE have reported that between-site heterogeneity arising from the use of different fMRI scanner types and experimental settings has an impact on the image properties of rs-fMRI data, and that this consequently impacts any rs-fMRI analysis (Nielsen et al., 2013; Castrillon et al., 2014).
- **Discriminative features:** *can we design new rs-fMRI features for better autism classification?* As pointed out above, powerful and complex ML methods such as RNN, GCN, and denoising autoencoders give similar top classification performance of less than 1% difference, while they all employ conventional brain *functional connectivity* (FC) features.

### 1.1. Domain adaptation

Domain adaptation methods operate on datasets from different sources with mismatched distributions to find a new latent space where the data is homogeneous, or source invariant (Pan & Yang, 2009; Weiss et al., 2016). In the context of this study, this corresponds to aligning the rs-fMRI data so that there is *independence* between the data and acquisition sites. Recently, Moradi et al. (2017) proposed a domain adaptation approach to correct site heterogeneity for the estimation of symptom severity in autism using data from four ABIDE sites. They first used partial least squares regression to identify a feature space where cortical thickness data extracted from structural MRI was independent from the four sites. Then they applied regression methods onto the site-adapted data to predict autism severity scores for each subject. Their severity score predictions were markedly better than those from models without domain adaptation. However, their study was limited to a small sample of 156 subjects from four of the 20 ABIDE sites and they did not tackle the classification problem. In contrast, our study focuses on the technical challenge of assessing and targeting the site heterogeneity in all 20 ABIDE sites to improve autism classification.

### 1.2. Functional connectivity

FC measures are important features in ASD classification. Two FC measures are widely used: 1) the Pearson correlation measures the coupling between pairs of regions of interest (ROIs), and 2) the more recent tangent embedding parameterisation of the covariance matrix proposed by Varoquaux et al. (2010) captures the FC differences between a single subject and a group. In this paper, we explore a new perspective: *for any two ROIs, are they functionally connected to other brain regions in the same way?* This inspires us to propose a new second-order FC measure that jointly considers the FC of individual ROIs.

### 1.3. Contributions

In this study, we analysed the rs-fMRI data of 1035 subjects from all 20 ABIDE sites to improve multi-site autism classification. We placed a specific focus on constructing new second-order FC measure and evaluating the impact of minimising their dependence on the acquisition sites for autism classification. The main contributions can be summarised as follows:

1. We proposed a new second-order FC measure, *Tangent Pearson (TP) embedding* to extract more discriminative features for multi-site autism classification. The proposed TP FC measure outperformed two commonly used FC measures on the whole. We also reported the neural patterns that are most informative for autism classification using this measure, for further biomarker analysis by neuroimaging researchers.
2. We assessed the statistical significance of the dependence between FC features and ABIDE acquisition sites, showing that across different feature representations, the dependence is significant at the 5% level. This provides a strong basis for designing models that correct for the between-site heterogeneity in the ABIDE dataset.
3. We applied a domain adaptation approach to minimise the dependence between ABIDE acquisition sites and FC features. Results demonstrated that minimising between-site heterogeneity leads to improvements in autism classification when combined with the TP measure and phenotypic information, yielding state of the art results.

The code for reproducing the experimental results of this study is publicly available at https://github.com/Mwizakunda/fMRI-site-adaptation.

## 2. Material and methods

Figure 1 gives an overview of the pipeline for studying domain adaptation on FC features. It shows the steps involved in specifying various models. Step 3 is the proposed domain adaptation step optional in the pipeline and it is used to extract site-invariant features from the FC data of step 2 in an *unsupervised* way. The impact of using such features can then be compared with models that do not use step 3. Likewise, the including of phenotypic information is optional.

**Figure 1:**
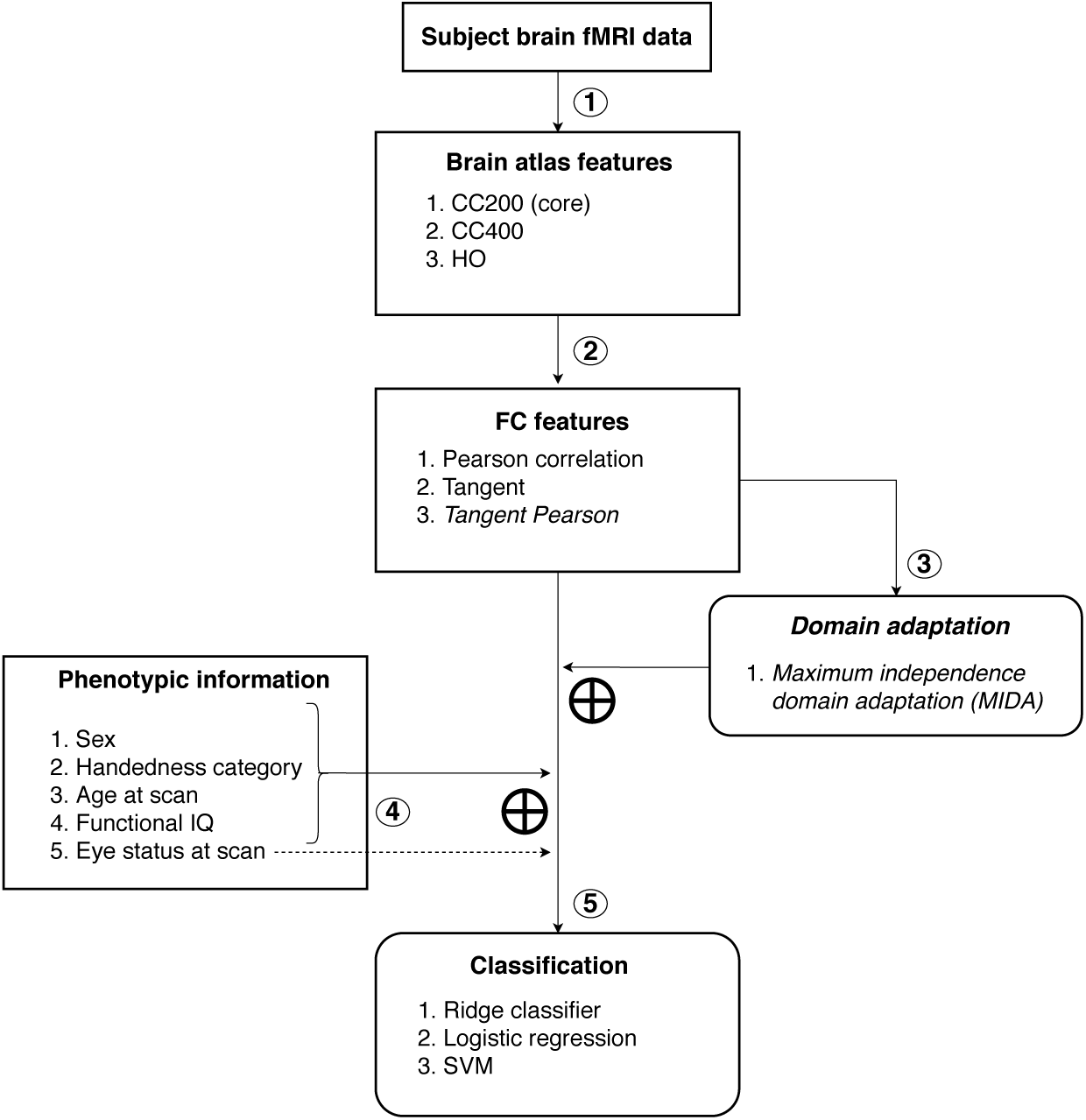
Overview of the pipeline for domain adaptation on functional connectivity (FC) features. An illustration of all possible models studied in this paper. Steps 1, 2 and 5 are compulsory, whilst 3 and 4 are optional. Step 3 applies maximum independence domain adaptation (MIDA) (Yan et al., 2017) to FC features and step 4 supplements subject feature vectors with phenotypic information. Phenotypes 1–5 are added for all phenotype-including models *not involving* MIDA. Phenotypes 1–4 are added for all phenotype-including models *involving* MIDA because the phenotype ‘eye status at scan’ is a site specific protocol. ‘**⊕**’ denotes concatenation.

### 2.1. ABIDE database: rs-fMRI and phenotypic data

This study focuses on the ABIDE database, which is composed of MRI and phenotypic data collected from 20 sites around the world. We included rs-fMRI and phenotypic data from 505 ASD and 530 TC individuals, yielding a sample of 1035 subjects. This sample of subjects is the same as that used in (Heinsfeld et al., 2018), which differs from the 871 subjects used in (Parisot et al., 2018; Abraham et al., 2017) due to their use of image quality control measures upon the full database. We opted for such a large sample to increase the likelihood of detecting site effects from individual sites, even though a larger sample presents the challenge of a greater level of heterogeneity expressed in subjects.

The ABIDE dataset provides a range of phenotypic information, including factors such as sex, age, full IQ (FIQ) test scores and handedness (left, right or ambidextrous). Table 1 gives a summary of each site with respect to key experimental protocols and phenotypic information. It shows that the type of fMRI scanner and length of individual fMRI scans varied across sites, giving rise to the apparent heterogeneity.

**Table 1:** Phenotypic and experimental variation across ABIDE sites. For quantitative variables, the standalone values represent the observed means, SD represents the standard deviation. M: male, F: female, R: right hand dominance.

| Site ID | SCANNER | DOMINANT<br>HAND | EYE<br>STATUS | SEX | AGE (SD) | FIQ (SD) | SCAN<br>TIME (SD) |
| --- | --- | --- | --- | --- | --- | --- | --- |
| CALTECH | SIEMENS Trio | R | Closed | M | 27.72 (10.45) | 111.16 (11.53) | 146.0 (0.0) |
| CMU | SIEMENS Verio | R | Closed | M | 26.59 (5.69) | 114.56 (10.54) | 273.26 (42.46) |
| KKI | Philips Achieva | R | Open | M | 10.01 (1.27) | 106.17 (15.0) | 148.5 (9.36) |
| LEUVEN_1 | Philips INTERA | R | Open | M | 22.59 (3.55) | 112.21 (13.03) | 246.0 (0.0) |
| LEUVEN_2 | Philips INTERA | R | Closed | M | 14.09 (1.38) | 100.0 (0.0) | 246.0 (0.0) |
| MAX_MUN | SIEMENS Verio | R | Closed | M | 25.31 (11.88) | 110.52 (11.73) | 140.62 (37.28) |
| NYU | SIEMENS Allegra | R | Open | M | 15.26 (6.57) | 110.51 (14.99) | 176.0 (0.0) |
| OHSU | SIEMENS Trio | R | Open | M | 10.71 (1.79) | 110.6 (16.81) | 78.0 (0.0) |
| OLIN | SIEMENS Allegra | R | Open | M | 16.59 (3.47) | 112.41 (17.01) | 206.0 (0.0) |
| PITT | SIEMENS Allegra | R | Closed | M | 18.94 (6.93) | 110.18 (12.24) | 196.0 (0.0) |
| SBL | Philips Intera | R | Closed | M | 34.37 (8.6) | 101.53 (6.15) | 196.0 (0.0) |
| SDSU | GE MR750 | R | Open | M | 14.41 (1.84) | 109.36 (13.77) | 176.0 (0.0) |
| STANFORD | GE Signa | R | Closed | M | 9.98 (1.59) | 111.41 (15.56) | 209.77 (30.02) |
| TRINITY | Philips Achieva | R | Closed | M | 16.96 (3.47) | 109.96 (13.75) | 146.0 (0.0) |
| UCLA_1 | SIEMENS Trio | R | Open | M | 13.19 (2.4) | 103.46 (12.02) | 116.0 (0.0) |
| UCLA_2 | SIEMENS Trio | R | Open | M | 12.49 (1.53) | 102.04 (14.92) | 116.0 (0.0) |
| UM_1 | GE Signa | R | Open | M | 13.4 (2.88) | 104.96 (14.11) | 296.0 (0.0) |
| UM_2 | GE Signa | R | Open | M | 16.01 (3.36) | 112.26 (10.85) | 296.0 (0.0) |
| USM | SIEMENS Trio | R | Open | M | 22.69 (8.34) | 105.23 (17.65) | 235.94 (0.47) |
| YALE | SIEMENS Trio | R | Open | M | 12.71 (2.88) | 99.77 (20.12) | 196.0 (0.0) |

To compare against the state-of-the-art (SOTA) methods (Parisot et al., 2018; Heinsfeld et al., 2018; Abraham et al., 2017), we used the same fMRI dataset from ABIDE^1^, which was processed by the Configurable Pipeline for the Analysis of Connectomes (CPAC) (Craddock et al., 2013).

### 2.2. Step 1: Brain atlas features

It is common for studies on rs-fMRI data to define brain regions of interest (ROIs) rather than operating on individual voxels. These ROIs represent the aggregation (e.g. averaging) of the rs-fMRI time series data of individual voxels, so that the number of ROIs is significantly less than the number of voxels.

We chose the Craddock 200 (CC200) brain atlas (Craddock et al., 2012) due to its robust performance in previous studies that used large subsets of the ABIDE dataset (>1000 subjects) (Dvornek et al., 2018; Heinsfeld et al., 2018; Parisot et al., 2018). CC200 has 200 ROIs derived from the clustering of spatially close voxels. We also considered two additional atlases to assess the impact of using a different brain parcellation: 1) Harvard Oxford (HO), a structural atlas with 110 ROIs based on anatomical landmarks from 40 sMRI scans (Makris et al., 2006), and 2) Craddock 400 (CC400), an atlas with 392 ROIs computed in a similar way to the CC200 atlas. For these three atlases, the representative time series of an ROI was derived by averaging the rs-fMRI time series of voxels associated with the ROI.

### 2.3. Step 2: Functional connectivity features

Instead of operating on the raw time series data of ROIs, FC features are usually extracted between pairs of ROIs based on the time series data. They estimate the fluctuating coupling of brain regions with respect to time, so that we can train predictive models to classify ASD and TC subjects based on differences in brain region coupling.

We first consider two SOTA FC measures as baselines: 1) the Fisher transformed *Pearson*’s correlation coefficient used in (Parisot et al., 2018), which gives a measure of coupling between pairs of ROIs by computing the correlation between their time series, and 2) the *tangent embedding* parameterisation of the covariance matrix proposed in (Varoquaux et al., 2010), which captures the deviation of each subject covariance matrix from the group mean covariance matrix and outperforms many other FC measures in the study by Abraham et al. (2017). These two FC measures have achieved SOTA autism classification performance on the ABIDE dataset.

#### Proposed second-order FC measure

As reviewed in Sec. 1, various ML methods including RNN and GCN have been applied on the above FC features for multi-site autism classification. However, the resulting top classification accuracies differ by less than 1%. This makes us question whether such simple FC measures can capture well the complexity of brain networks. Thus, it motivates us to go a step further than the two standard FC measures and construct a new *second-order* FC measure in the following.

The Pearson correlation coefficient gives a measure of the coupling between pairs of ROIs irrespective of any other ROIs. We explore a new direction to also quantify the relationship that two ROIs have with respect to all other ROIs. That is, we want to examine the following question: *for any two ROIs, are they functionally connected to other brain regions in the same way?*

Given a set of *R* ROIs and a corresponding FC matrix, **M** (*R* × *R*), each row of **M, M**_*i*_, *i* ∈ {1, …, *R*}, gives the measure of FC between region *i* and all other regions. So given any two regions *i* and *j*, the *second-order* measure of interest can therefore be computed by measuring the similarity between **M**_*i*_ and **M**_*j*_. We propose to capture this second-order measure by firstly computing the Pearson correlation coefficient between pairs of ROI time series, and then computing the covariance of the resulting connectivity profiles from all regions (e.g. **M**_*i*_ and **M**_*j*_). The Pearson correlation coefficient is used at the first step because it is powerful in capturing first-order FC measures, while the covariance is used as the second-order similarity measure so that we can leverage the *tangent embedding* connectivity for a group-based comparison of each subject FC.

We name this proposed measure as the *Tangent Pearson embedding FC measure*, or simply *Tangent Pearson* (TP) because it can be seen as combination of the Pearson and tangent embedding FC measures with a *simple implementation*: replacing the covariance matrices used in the original tangent embedding method with the proposed correlation-based second-order FC matrices so that the deviation of each subject from the group is computed from the group mean second-order FC matrix.

The resulting FC matrices computed for each subject using these three FC measures are symmetric. Therefore, it is sufficient to keep only the upper/lower triangular parts. We chose to keep the upper triangular parts and also discarded the main diagonal FC values since they represented the self connectivity of an ROI with itself, which is redundant information. The remaining upper triangular values were flattened into a one-dimensional vector and used as FC features for each subject in all subsequent analyses.

### 2.4. Statistical test of independence

Despite the aggregation of voxels’ rs-fMRI time series and the estimation of FC features, acquisition site effects have been observed in the results of studies using these FC features. For example, Plitt et al. (2015) identified large univariate differences in FC strength between three of the sites they investigated (NYU, UCLA 1 and USM), showing the persistence of site effects into FC features. Parisot et al. (2018) and Nielsen et al. (2013) highlighted that significantly higher accuracies can be obtained on single-site studies in comparison to multi-site studies, however, this is also due to the reduced heterogeneity of both ASD and TC subjects in single-site studies. Before dealing with the dependence between ABIDE acquisition sites and FC features, we aim to first assess the statistical significance of this dependence. This assessment would provide a measure of the influence of site effects on FC features, and whether the effect is large enough to merit being accounted for during the modelling of ASD classifiers on the ABIDE dataset.

To conduct this statistical test, we employed the Hilbert-Schmidt independence criterion (HSIC), an empirical kernel-based statistical independence measure. It is superior to other kernel-based independence measures due to being simpler, converging faster and having a low sample bias with respect to the sample size (Gretton et al., 2005, 2008). We firstly used the HSIC to measure the statistical dependence between ABIDE sites and FC features. We then evaluated the statistical significance of such dependence using a hypothesis test derived for the HSIC by Gretton et al. (2008).

#### 2.4.1. Hilbert-Schmidt independence criterion

Given two multivariate random variables **X** and **Y** with associated probability distributions *P*_**X**,**Y**_, *P*_**X**_ and *P*_**Y**_, the HSIC provides a non-parametric way of measuring their statistical dependence. The HSIC gives a measure of zero if they are independent, and a value greater than zero otherwise. The larger the value of the HSIC, the stronger the dependence between them. Empirically, given *n* realisations for the random variables **X** = {**x**_*i*_} and **Y** = {**y**_*i*_}, the HSIC between **X** and **Y**, denoted by *ρ*_*h*_(**X, Y**), is given by Gretton et al. (2005)

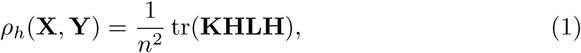

where **K, H, L** ∈ ℝ^*n*×*n*^, **K**_*i,j*_ = *k*_*x*_ (**x**_*i*_, **x**_*j*_) and **L**_*i,j*_ = *k*_*y*_ (**y**_*i*_, **y**_*j*_). *k*_*x*_(·) and *k*_*y*_(·) are two kernel functions, e.g. linear, polynomial, or radial basis function (RBF). 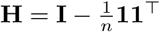 is a centering matrix and tr(·) is the trace function.

For our problem, we define **X** to be the random variable corresponding to FC features so that **x**_*i*_ is a single subject FC features. We define **Y** to be the random variable corresponding to the acquisition site, with **y**_*i*_ ∈ ℝ^20^ a one-hot encoding of each site (more detail in Sec. 2.5). For the kernel functions, a linear kernel was used for *k*_*y*_(·) due to the theoretical results in Zhou et al. (2020), where a correlation between HSIC and distribution divergence measure is guaranteed by linear kernel. RBF kernel was used for *k*_*x*_(·) in order to model non-linear dependence between FC features and sites. We set the width parameter of the RBF kernel, *σ*, with the median distance between FC features.

#### 2.4.2. Measure of significance

Gretton et al. (2008) proposed to measure the statistical significance of an HSIC estimate based on a hypothesis test of independence for two random variables **X** and **Y** using the HSIC estimate *ρ*_*h*_(**X, Y**) as a test statistic. The test considers a null hypothesis of independence ℋ_0_ : *P*_**X**,**Y**_ = *P*_**X**_*P*_**Y**_ against an alternative hypothesis ℋ_*A*_ : *P*_**X**,**Y**_ ≠ *P*_**X**_*P*_**Y**_. Evidence for the acceptance of the null hypothesis is obtained by comparing the test statistic *ρ*_*h*_(**X, Y**) against a threshold *T* such that if *ρ*_*h*_(**X, Y**) ≤ *T*, the null hypothesis can be accepted. In other words, w.r.t. *T*, the HSIC estimate is sufficiently close to zero for independence between **X** and **Y** to be accepted. In (Gretton et al., 2008), this threshold is set to be the 1−*λ* (*λ* ∈ [0, 1]) quantile of the null distribution for the test statistic, which is approximated by a two-parameter Gamma distribution.

### 2.5. Step 3: Domain adaptation

To target the inter-site heterogeneity, we applied a recently proposed multisource domain adaptation approach, maximum independence domain adaptation (MIDA) (Yan et al., 2017). Since we hypothesise that there exists inter-site difference in the FC features of subjects from different sites, the aim is to extract new features from FC data that are no longer site dependent, and are consequently more performant for classification.

MIDA obtains these site-invariant features in an unsupervised manner, utilising the empirical HSIC introduced in Sec. 2.4.1 as a measure of dependence. Given a multivariate random variable **S** for the acquisition site, a projection map parameterised by **W**, *ϕ*_**W**_(·), and a random variable **X** for the subject FC features, MIDA aims to learn **W** such that the empirical HSIC between the random variables *ϕ*_**W**_(**X**) and **S** is close to zero. In other words, in the space *ϕ*_**W**_(**X**), the data are independent of their respective acquisition sites. More specifically, let **x**_*i*_ ∈ ℝ^*k*^ be the FC features of subject *i* where *i* = 1, …, 1035 and *k* is the dimensionality of FC features. MIDA learns a low-dimensional feature **z**_*i*_ ∈ ℝ^*h*^, where *h* ≤ *n*.

To apply MIDA, we need to construct a site feature vector **d**_*i*_ ∈ ℝ^*v*^ for each subject to encode information about their respective acquisition site. The simplest is the site label information (e.g. CALTECH, CMU, KKI), which can be encoded with a one-hot scheme:

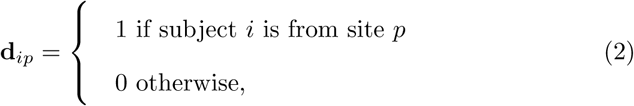

where **d**_*ip*_ is the *p*th entry of **d**_*i*_ and the number of sites *v* = 20. We concatenated site and FC feature vectors to encode each subject’s FC features with site information: **x**_*i*_ = [**x**_*i*_, **d**_*i*_]. Then we learned a mapping **W** ∈ ℝ^*n*×*h*^ to project the augmented features to a new subspace where the new feature representations for all subjects are minimally dependent on the site:

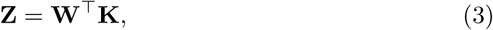

where **K** ∈ ℝ^*n*×*n*^, **K**_*i,j*_ = *k*_*x*_ (**x**_*i*_, **x**_*j*_) and each column of **Z** ∈ ℝ^*h*×*n*^, **z**_*i*_, is the new feature representation of the augmented features. Next, we formulated the objective function by maximising the preserved data variance while minimising the statistical dependence on the site, i.e.

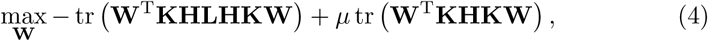

where *µ* > 0 is a hyperparameter governing the emphasis of variance preservation against the level of independence achieved between the projected features **Z** and the site features. The solution can be found by forming **W** from the eigenvectors of the matrix **K** (−**HLH** + *µ***H**) **K** corresponding to the *h* largest eigenvalues (Yan et al., 2017).

The hyperparameters are *µ, h* and the kernel functions *k*_*x*_(·), *k*_*d*_(·). We used a linear kernel for *k*_*d*_(·) and the RBF kernel for *k*_*x*_(·) with the width parameter set to the median distance between FC features as in Sec. 2.4.1 and optimised *µ* and *h* via a grid-search scheme detailed in Sec. 2.8.

### 2.6. Step 4: Incorporating phenotypic information

The phenotypic information included in the ABIDE dataset for each subject is extensive. Including these features when training a classifier for ASD has been shown to be beneficial (Parikh et al., 2019). Several studies on autism have observed sex and age-related differences between ASD and TC subjects. Werling & Geschwind (2013) identified sex-differential genetic and hormonal factors that supported the observation that females are typically less frequently affected by ASD than males. In (Kana et al., 2015), age matched ASD and TC children were found to have differences in FC. In fact, between ASD patients, FC differences have also been observed with respect to age (Uddin et al., 2013; Supekar et al., 2013). However, these findings are not yet conclusive (Müller et al., 2011).

Recent studies (Dvornek et al., 2018; Parisot et al., 2018) on the ABIDE dataset improved ASD classification accuracy by leveraging phenotypical information. We proceeded in a similar way and assessed the impact of including phenotypes in ASD classifiers. We considered only sex, age, full IQ (FIQ), handedness and eye status at scan since the majority of subjects had such information present. For each of the categorical variables (handedness, sex and eye status at scan), a one-hot encoding scheme was used to construct phenotype features for each subject and *concatenated* with other features before feeding into the classifier. For subjects with missing values for FIQ and handedness, we used the same imputation method used in (Dvornek et al., 2018): 1) *Handedness* – right hand dominance was assigned since most people are right-handed; 2) *FIQ* – a score of 100 was assigned which is considered the average IQ score.

### 2.7. Step 5: Classification

The impact of removing site effects can be assessed by comparing using the “raw” FC features (Sec. 2.3) against site-invariant features derived from MIDA as inputs to a classifier. Here we prefer linear models (over deep learning models) to make isolating the impact of domain adaptation less complex and allow for a greater degree of interpretability (Bishop, 2006), e.g. by visualising the model coefficients to identify any functional differences between ASD and TC subjects.

We chose three standard linear classifiers: ridge classifier (ridge regression with binary target values), logistic regression (LR), and support vector machines (SVM) from the Scikit-learn library (Pedregosa et al., 2011). For all models, the hyperparameter values were selected via a grid-search scheme detailed in Sec. 2.8.

### 2.8. Experimental setup

We designed the experiments with three objectives:

- To test the statistical dependence between acquisition sites and FC features.
- To assess the impact of the proposed second-order FC measure and site-dependence minimisation on autism classification.
- To extract biomarkers from the trained ML models to distinguish between ASD and TC.

#### Algorithm setting

We tested various models that can be constructed from the pipeline in Fig. 1, including both existing and proposed ones. We used the popular CC200 as the default atlas. Table 2 shows the hyperparameters for grid search. We evaluated all possible combinations of three values for each on the training data via five-fold cross validation (CV) to find the best setting.

**Table 2:** *Hyperparameter setting for the ML models. h*: the number of eigenvectors in MIDA; *µ*: the weighting of variance maximisation in MIDA; *C*: the *l*_2_ regularisation coefficient for the three classifiers (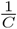 for logistic regression and SVM).

| ML Method | Hyperparameter 1 | Hyperparameter 2 |
| --- | --- | --- |
| MIDA | $h = 50, 150, 300$ | $\mu = 0.5, 0.75, 1.0$ |
| Ridge classifier | $C = 0.25, 0.5, 0.75$ | NA |
| Logistic regression | $C = 1, 5, 10$ | NA |
| SVM | $C = 1, 5, 10$ | NA |

#### Statistical test of independence

We assessed the independence between sites and FC features using the statistical test in Sec. 2.4. We set the significance level *λ* := 0.05 so that the probability of rejecting the null hypothesis when it is true is 0.01. In particular, given an observed HSIC estimate from the data, *ρ*_*h*_(**X, Y**), the null hypothesis was set to be rejected at the 5% level if *ρ*_*h*_(**X, Y**) > *t*_1−*λ*_ with *t*_1−*λ*_ being the 95% quantile of the estimated Gamma distribution. The kernel functions *k*_*y*_(·) and *k*_*x*_(·) were defined according to the empirical HSIC estimate detailed in Sec. 2.4.1.

We followed the terminology in (Abraham et al., 2017) to consider the *intrasite* and *inter-site* prediction.

### Intra-site prediction

This is the most commonly used setting (Parisot et al., 2018; Abraham et al., 2017; Heinsfeld et al., 2018), where the data from all 20 sites are mixed to form training/test sets with the same proportion of ASD/TC for stratified 10-fold CV. We compare MIDA-based models with those without using MIDA as baseline *raw* models to assess the impact of MIDA. We report the average accuracy and Area Under the Receiver Operating Characteristics (AUROC) over the 10 folds for each model. We also studied the impact of adding phenotypic features as in the studies by Parisot et al. (2018) and Dvornek et al. (2018). For the MIDA-based models, ‘eye status at scan’ was not included as a phenotypic measure since it represents a site-specific protocol. Additionally, we evaluated impact of brain atlas by validating MIDA-based models on the CC200, CC400, and Harvard-Oxford (HO) atlases.

### Inter-site prediction

This setting uses data from one individual site as testing data while training on the data from all 19 remaining sites to study the generalisation performance to sites unseen in training. This is more challenging than the intra-site setting. The average accuracy/AUROC over the 20 sites will be reported. However, since each site has a different sample size, we computed the average by weighting the contribution of each site by sample size. Specifically, we measured the average accuracy by simply counting the total number of correct predictions across all sites from 20 runs and dividing by the total sample size. For AUROC, we similarly computed an overall measure across all sites. We also assessed the influence of phenotypic information as above.

### Comparison with other studies

We compared our proposed models with those in recent studies (Parisot et al., 2018; Heinsfeld et al., 2018; Abraham et al., 2017) in both intra-site and inter-site settings. For inter-site setting, Abraham et al. (2017) and Heinsfeld et al. (2018) computed the average accuracy without weighting by sample size so we will report both the weighted and unweighted average accuracies for fair comparison. Moreover, we studied two sample sizes, 871 and 1035, which have been studied in previous studies. For completeness, we also applied the SOTA method by Parisot et al. (2018) to the larger sample size of 1035 using their implementation^2^. To account for different stratifications in 10-fold CV between our study and (Parisot et al., 2018), we used *the same stratification* as in (Parisot et al., 2018) but also evaluated both methods using five different stratifications to reduce the influence of the stratification. In addition, we conducted two-sample Welch *t*-tests to assess the statistical significance of performance improvements in intra-site evaluation and will report the corresponding *p*-values. For the inter-site setting, since we compute summary metrics without averaging and thus only have single representative values, we did not conduct *t*-tests to assess significance.

#### Biomarker extraction

Linear classifiers evaluate **w**^┬^**x** to make a decision, where the model weight **w** is the learned decision hyperplane and **x** is the feature vector. In our context, the coefficients of **x** in **w** indicate the informativeness of each feature in **x** in classifying a subject as ASD or TC. If **x** is the (flattened) FC features, the values of the parameters in **w** will indicate which pairs of ROIs are important to distinguish ASD and TC. Positive and negative elements of **w** indicate ROI-ROI connections that are informative for ASD and TC, respectively. The larger the absolute value of the coefficient of a given ROI-ROI FC, the more informative this FC. We analysed all the ten **w**s from the 10-fold CV in the intra-site setting to extract ROI-ROI connections that are consistently the most informative for classification. Specifically, we identified the top 50 largest weights in absolute value for each fold individually and then ranked ROI-ROI FC connections for consistent occurrence in the top 50 features across all 10 folds, so that the most informative connections would be present in the top 50 features across all 10 folds. Where two ROIs had the same number of occurrence across folds (e.g. 7 times), we sorted the ranking by the absolute value of their respective coefficient averaged across folds.

## 3. Results

### 3.1. Statistical test of independence

Table 3 shows the statistical test of independence between FC features and sites. From the table, irrespective of the FC, the null hypothesis of independence between FC features and sites can be rejected with 95% confidence because the sample HSIC estimate exceeds the threshold. Also, the *p*-values are extremely small (< 10^−5^), giving greater evidence for rejecting the null hypothesis.

**Table 3:**
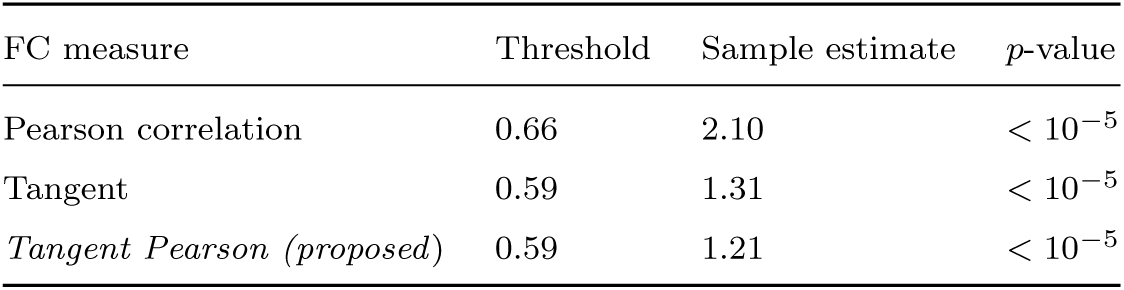
The statistical test of independence. The threshold value gives the 95% quantile of the estimated Gamma distribution. The sample estimate gives the sample HSIC estimate, with the corresponding *p*-value computed. The difference in the threshold values arises from the different Gamma distribution approximations for different FC measures.

| FC measure | Threshold | Sample estimate | $p$-value |
| --- | --- | --- | --- |
| Pearson correlation | 0.66 | 2.10 | $< 10^{-5}$ |
| Tangent | 0.59 | 1.31 | $< 10^{-5}$ |
| *Tangent Pearson (proposed)* | 0.59 | 1.21 | $< 10^{-5}$ |

Next, we show the effect of domain-independent adapation by visualising features in a 2-D space. We applied Principal Component Analysis (PCA) to reduce the dimensionality of FC features to 50, and then employed *t*-distributed Stochastic Neighbour Embedding (*t*-SNE) (Maaten & Hinton, 2008) to project them to a 2-D space for visualisation. Figure 2 shows the *t*-SNE projections of the proposed tangent Pearson FC features w.r.t. acquisition sites for both with/without site adaptation. In subplot (a) without adaptation, site-specific clusters can be identified (SDSU, SBL, TRINITY, etc.) while in subplot (b) with adaptation, there is a reduction of association between FC features and acquisition site. This further illustrates the site specificity of the ABIDE data.

**Figure 2:**
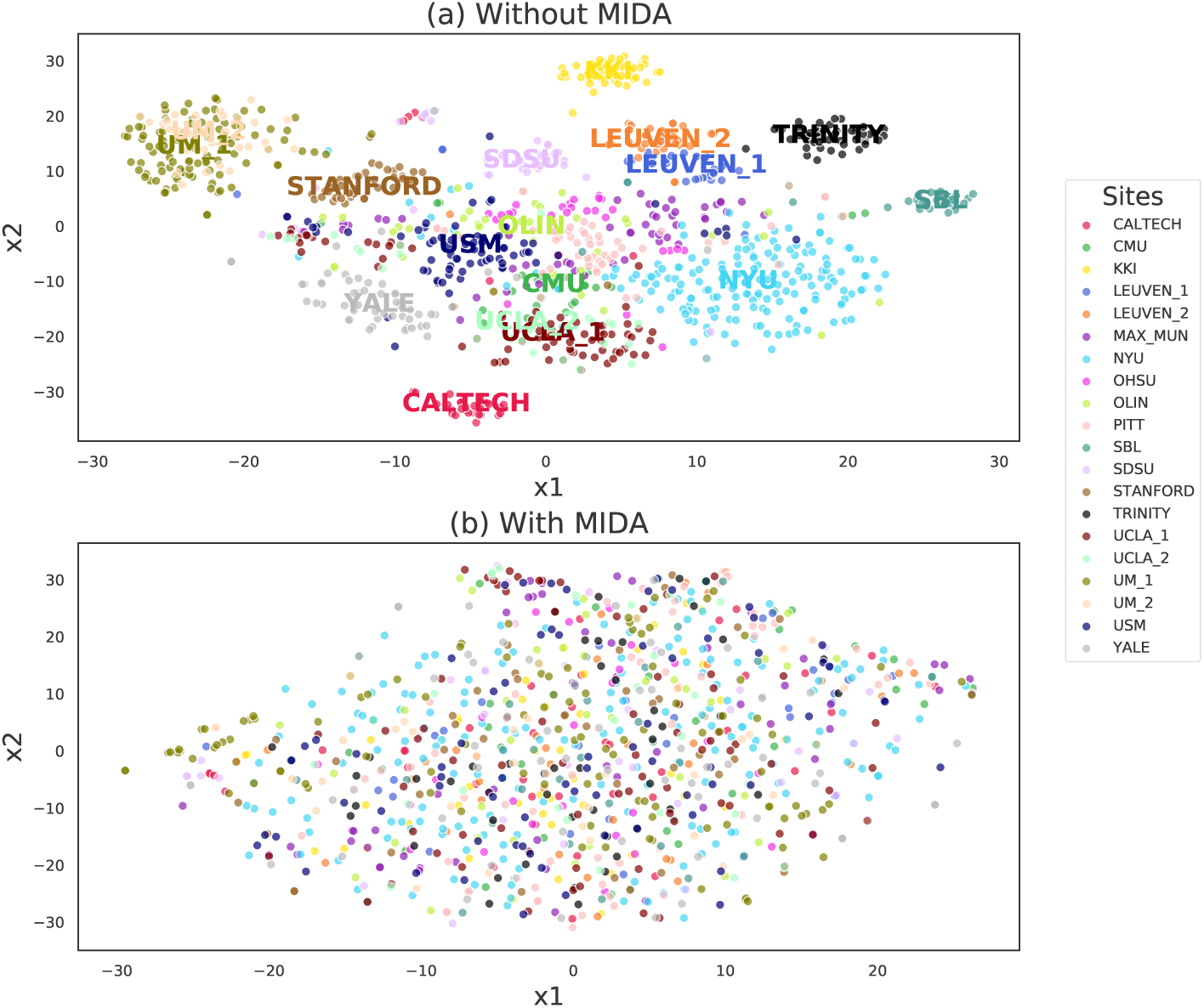
The effect of domain-independent adaptation. A 2-D *t*-SNE projection of CC200-generated tangent Pearson FC features with (a) no site adaptation, (b) site adaptation by MIDA, using the scikit-learn *t*-SNE implementation with perplexity 30 and learning rate 10. In (a), key identifiable clusters are labelled while the other site labels (OHSU, MAX_MUN, and PITT) are omitted since their clusters are not well defined. In (b), site labels are omitted because MIDA removed the association between features and sites.

### 3.2. Intra-site prediction

Figure 3 shows the intra-site prediction results. We analyse the results with focus on the impact of (a) FC features, (b) MIDA, and then (c) phenotypic features, on the classification performance.

**Figure 3:**
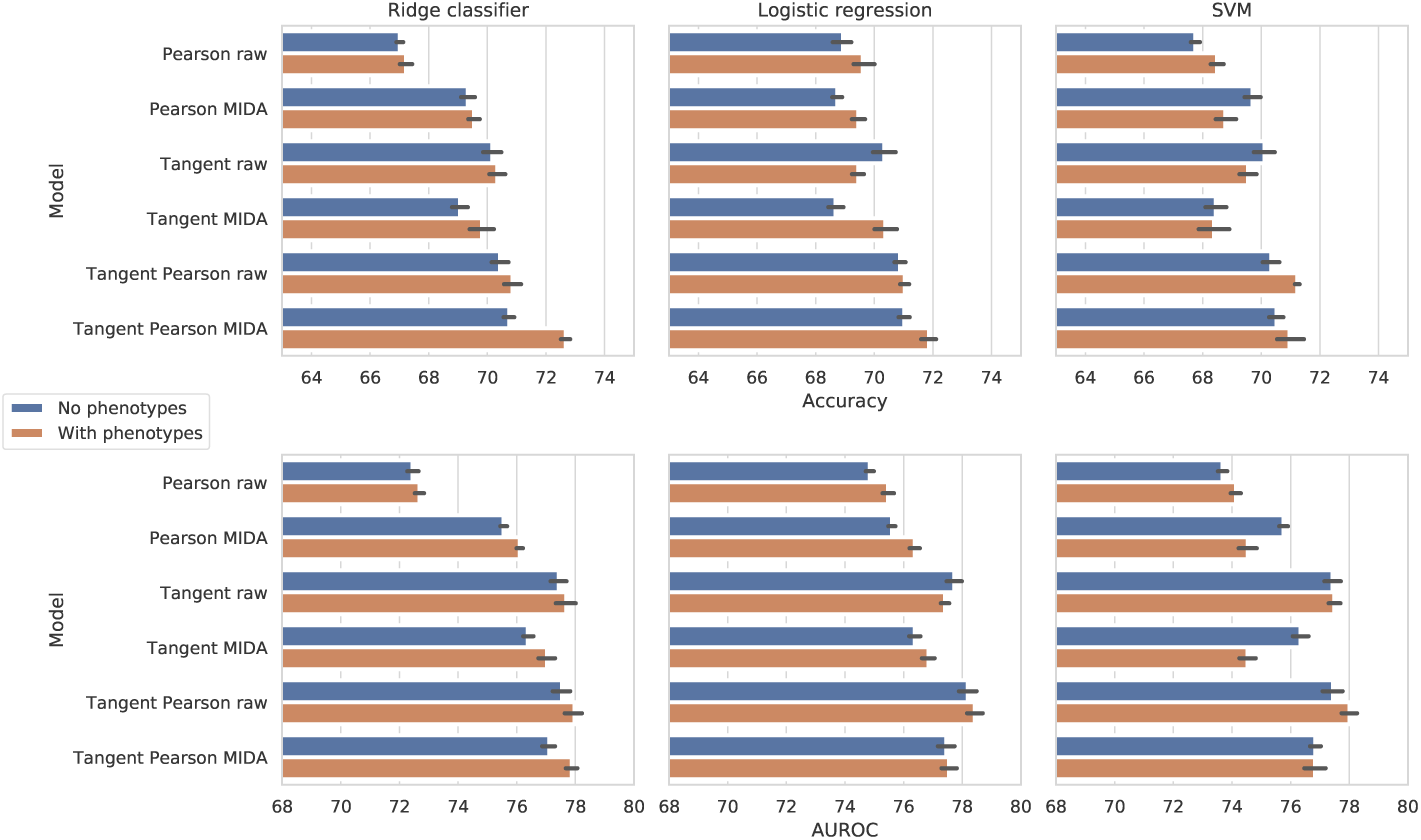
Intra-site prediction with 5×10-fold CV. A comparison of the effect of site adaptation (raw baseline vs MIDA), classifier (ridge, logistic, SVM), FC measure (Pearson, tangent embedding, proposed tangent Pearson), and the inclusion of phenotypic information, on the average accuracy and AUROC.

a. *Impact of FC measures*. For the baseline models (raw, without MIDA and phenotypes), the performance of the three FC measures is very stable across classifiers and CV splitting settings. For each classifier, the highest accuracy was consistently obtained by the proposed second-order FC measure, TP embedding. In contrast, the Pearson correlation gave the lowest accuracy for all classifiers.
b. *Impact of MIDA*. The effectiveness of MIDA seems to be sensitive to FC measure. Applying MIDA to Pearson features led to improvement in both accuracy and AUROC. In contrast, applying MIDA to tangent features led to worse results. Applying MIDA to TP embedding led to better accuracy but worse AUROC across different classifiers. A possible explanation for the performance drop is that there could be some loss of ASD/TC-specific information when learning site-invariant features using the tangent FC with MIDA, so that some subjects are not properly represented for classification. In Sec. 4, we will discuss further studies that can potentially reduce such unwanted effects.
c. *Impact of phenotypic features*. In most cases, adding the phenotypic features has improved the classification performance. The best accuracy is 72.7%, which is obtained using TP+MIDA with phenotypes. The positive impact of phenotypic information on autism classification observed here is consistent with the findings in previous studies (Parisot et al., 2018; Dvornek et al., 2018).

### 3.3. Impact of brain atlas

Table 4 compares the intra-site results of MIDA with tangent Pearson FC and phenotypic features (TP MIDA) on three different brain atlases: CC200, CC400, and HO. On the whole, there is no significant difference between different atlases, despite using HO atlas leads to a relative lower accuracy and AUROC for Ridge classifier. Thus, CC200 is a good choice for our MIDA-based models, as in other methods in the literature.

**Table 4:** Impact of brain atlas. MIDA with Tangent Pearson (TP) FC and phenotypic features using three different brain atlases in intra-site prediction. The 5*×*10-fold CV standard deviations (SD) over five different random seeds are in parentheses. ACC: accuracy.

|  | Ridge classifier |  | Logistic regression |  | SVM |  |
| --- | --- | --- | --- | --- | --- | --- |
| Atlas | ACC (SD) | AUROC (SD) | ACC (SD) | AUROC (SD) | ACC (SD) | AUROC (SD) |
| CC200 | 72.7 (0.3) | 77.9 (0.4) | 71.9 (0.6) | 77.6 (0.5) | 71.0 (1.0) | 76.8 (0.8) |
| CC400 | 72.2 (0.3) | 78.3 (0.5) | 71.4 (0.4) | 78.0 (0.6) | 71.1 (0.9) | 76.9 (1.3) |
| HO | 71.5 (1.0) | 77.0 (0.6) | 71.6 (1.0) | 77.1 (0.4) | 71.0 (0.5) | 77.5 (0.5) |

### 3.4. Inter-site prediction

Similarly, Figure 4 shows the inter-site results and we perform similar analyses as in the intra-site setting below.

**Figure 4:**
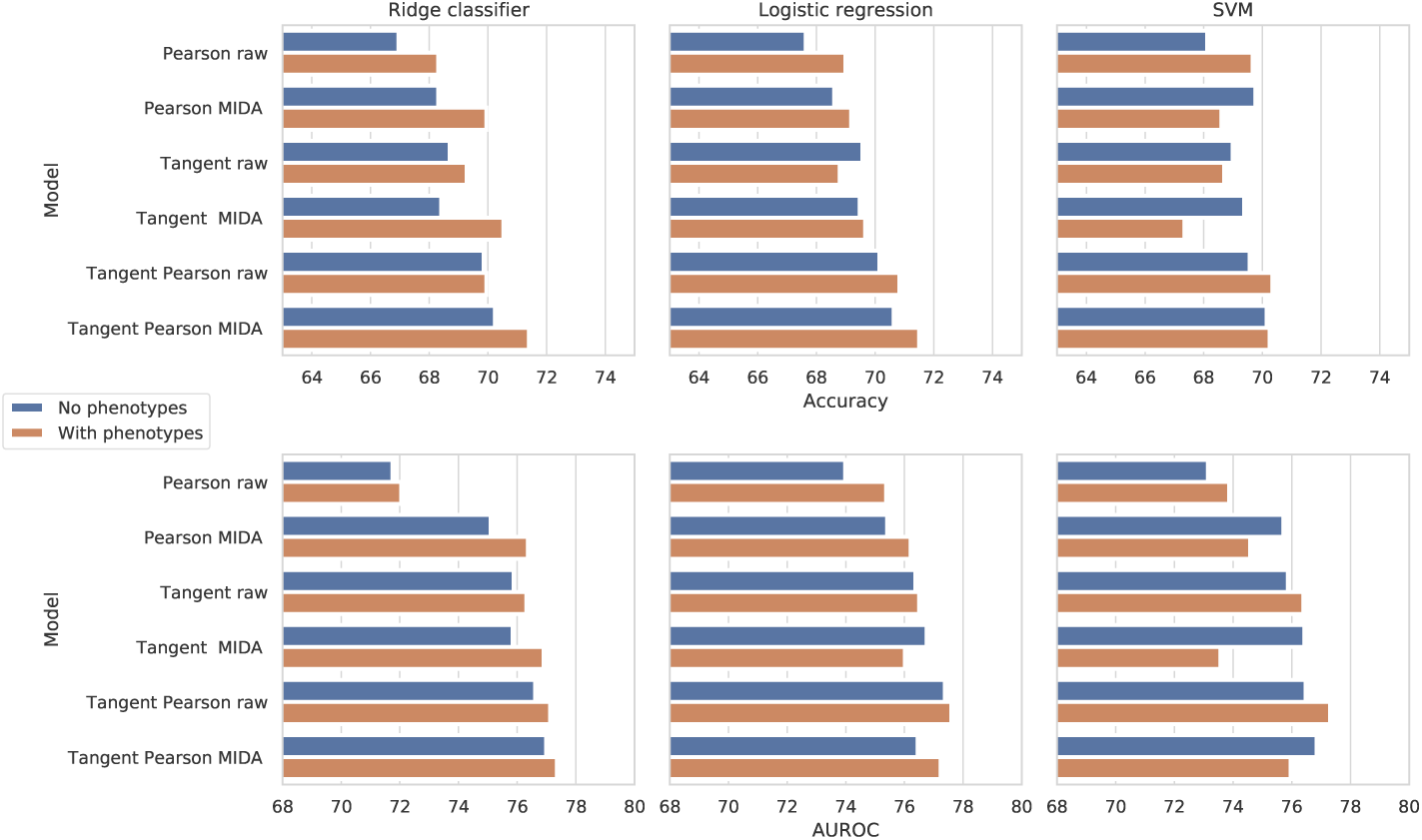
Inter-site prediction (20 runs). A comparison of the effect of site adaptation (raw baseline vs MIDA), classifier (ridge, logistic, SVM), FC measure (Pearson, tangent embedding, proposed tangent Pearson), and the inclusion of phenotypic information, on the weighted leave-one-site-out CV accuracy and AUROC.

#### (a) Impact of FC features

As with the intra-site setting, the tangent-based models achieve a dominant performance over the Pearson correlation-based ones. For the ridge and logistic regression classification models, the tangent Pearson measure achieves the highest accuracies of 69.9% (AUROC: 76.6%) and 70.1% (AUROC: 77.4%), respectively. The tangent measure achieves the highest accuracy of 69.6% (AUROC: 76.4%) with logistic regression. The Pearson correlation measure achieves its highest accuracy of 68.1% (AUROC: 73.1%) with SVM.

#### (b) Impact of MIDA

MIDA has an overall positive influence in this setting, with a reduced performance only for the tangent FC. For Pearson correlation and tangent Pearson measures, the use of MIDA has increased the accuracy by 0.63% (AUROC: 0.90%) over baseline equivalents, averaged across the three classifiers. The tangent Pearson measure achieves the highest accuracy of 70.6% (AUROC: 76.4%) with MIDA and logistic regression, while its non-MIDA equivalent achieves an accuracy of 70.1% (AUROC: 76.4%). Applying MIDA to tangent FC features leads to no significant difference in accuracy/AUROC w.r.t. baseline models across all classifiers.

#### (c) Impact of phenotypic features

Adding phenotypic features in inter-site prediction also helped improving the accuracy/AUROC of the baseline and MIDA models in most cases, e.g. an increases in accuracy by 1.05% (AU-ROC: 0.54%) on average over the three classifiers w.r.t. no phenotypic features. Overall, the highest inter-site accuracy of 71.4% (AUROC: 77.4%) is obtained with the phenotypic features, the application of MIDA to the tangent Pearson FC, and the ridge classifier. Its non-phenotype equivalent has a lower accuracy of 70.2% (AUROC: 77.0%).

#### (d) Factors for site accuracy variation

From the inter-site results above, MIDA-based models obtained better performance. However, the individual classification performance on each site varies a lot. Here, we investigate the following two potential factors that may affect the performance on an individual site:

1. *Mean length of rs-fMRI scan time at each site*. Intuitively, we expect that having a longer experimental scan time increases the ability to detect differences between ASD and TC subjects.
2. *Number of samples collected from site*. Sites with low sample sizes may be under-represented in the ABIDE database and may be significantly different in distribution from the other sites.

Figure 5 studies the correlations between site accuracy and site scan time or sample size from the results obtained by the model with tangent Pearson, MIDA, and ridge classifier but without phenotypic features. For scan time, a slight positive correlation between the mean scan time and the site accuracy can be identified (the left panel of Fig. 5). An interesting finding is that the lowest scoring site (OHSU) has the lowest mean scan time (78 seconds) in comparison to the other sites (all exceeding 116 seconds). Thus, it may be difficult to capture the difference between ASD and TC effectively in such a short period of time. Longer scan time has the potential of removing noise from rs-fMRI data and helping better capture signals differentiating autism or neurotypical effects. For site sample size, no conclusive relationship can be found with the observed site accuracy (the right panel of Fig. 5).

**Figure 5:**
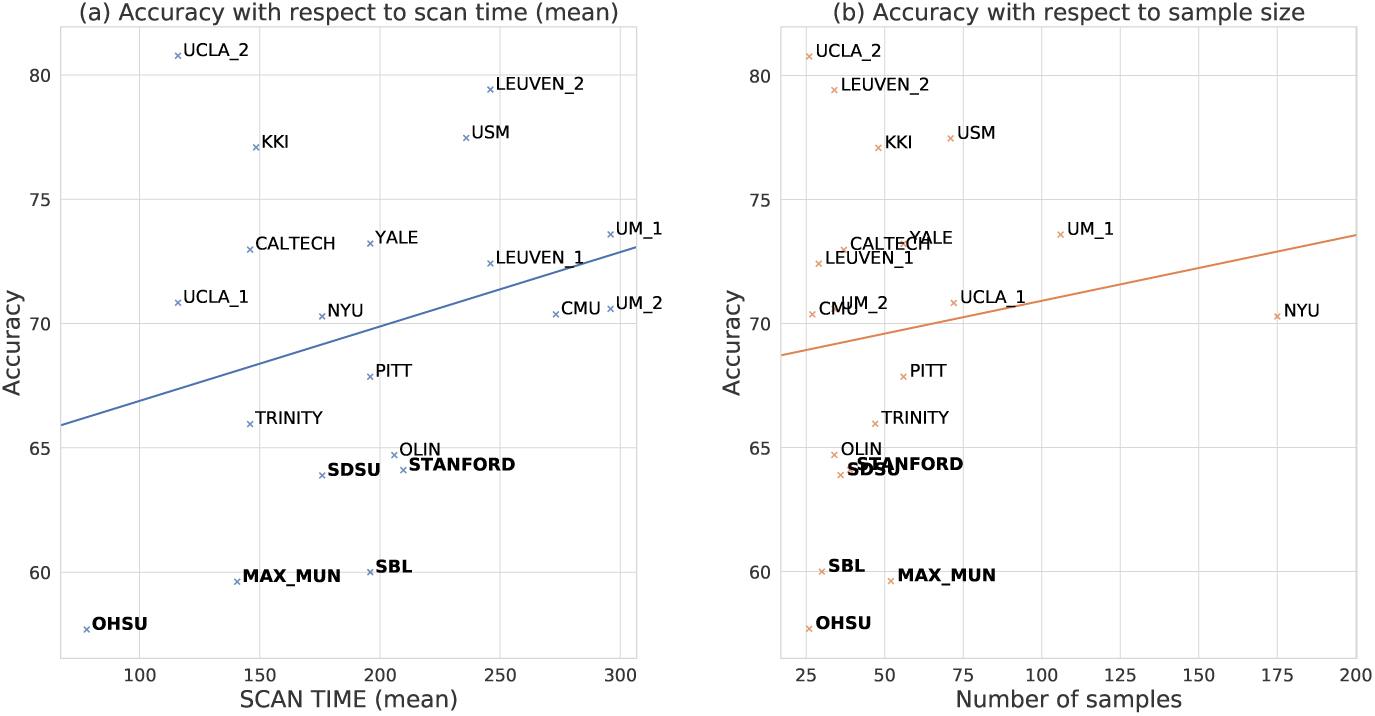
Potential factors for site accuracy variation. The sites are visualised w.r.t. the accuracy obtained by the tangent Pearson MIDA ridge classifier without phenotypic features.

### 3.5. Comparison with other studies

We evaluated our top performing models against three SOTA methods for both the intra-site and inter-site settings below.

#### Intra-site comparison

Table 5 reports the intra-site results. In 10-fold CV, our TP MIDA ridge (with phenotypic features) on a sample size of 1035 outperforms the SOTA methods in both accuracy and AUROC with scores of 73.0% and 78.0%, respectively. With respect to the accuracy of the SOTA (Parisot et al., 2018) on 871 and 1035 subjects, this is an increase of 2.6% (*p* = 0.09) and 4.8% (*p* < 10^−2^), respectively. In 5×10-fold CV, TP MIDA ridge achieves an increase of 4.8% (*p* < 10^−2^) and 4.1% (*p* < 10^−2^) in accuracy over (Parisot et al., 2018) on 871 and 1035 subjects, respectively.

**Table 5:** Intra-site comparison to other studies. ‘-’ indicates that the metric was not provided in the original study and we did not implement ourselves. The sample size is 871 if quality control (QC) was performed, and 1035 otherwise. We firstly report the results obtained using the same split setting of 10-fold CV in (Parisot et al., 2018). Then we show the results obtained under 5*×*10-fold CV with four more CV splittings generated by using different random seeds. Standard deviations for 10-fold CV were computed over 10 different partitions and those for 5*×*10-fold CV were computed over five different random seeds. Results of (Heinsfeld et al., 2018; Abraham et al., 2017) are cited from the original paper.

| Model | 10-fold CV |  |  | 5×10-fold CV |  |
| --- | --- | --- | --- | --- | --- |
|  | QC | ACC (SD) | AUROC (SD) | ACC (SD) | AUROC (SD) |
| TP MIDA | ✗ | 73.0 (3.9) | 78.0 (4.7) | 72.7 (0.3) | 77.9 (0.4) |
| TP raw | ✗ | 71.6 (2.7) | 78.2 (3.6) | 70.9 (0.6) | 78.0 (0.7) |
| Parisot et al. (2018) | ✗ | 68.2 (3.7) | 75.2 (3.8) | 68.6 (0.3) | 75.2 (0.5) |
| Heinsfeld et al. (2018) | ✗ | 70.0 (-) | - (-) | - (-) | - (-) |
| TP MIDA | ✓ | 70.0 (7.6) | 75.5 (7.3) | 69.7 (0.5) | 75.9 (0.5) |
| TP raw | ✓ | 69.2 (7.5) | 75.1 (7.1) | 70.1 (0.8) | 76.5 (0.9) |
| Parisot et al. (2018) | ✓ | 70.4 (3.9) | 75.0 (4.6) | 67.9 (1.3) | 73.3 (0.9) |
| Abraham et al. (2017) | ✓ | 66.9 (2.7) | - (-) | - (-) | - (-) |

When more training examples were included (using all 1035 samples without QC), there is a relative significant improvement obtained by TP MIDA while the change is small for non-adaptation models, i.e. TP raw and Parisot et al. (2018). The inter-site results in Table 6 has similar observations. We will discuss the relationship between those “poor quality” samples and domain adaptation effectiveness in Sec. 4.

**Table 6:**
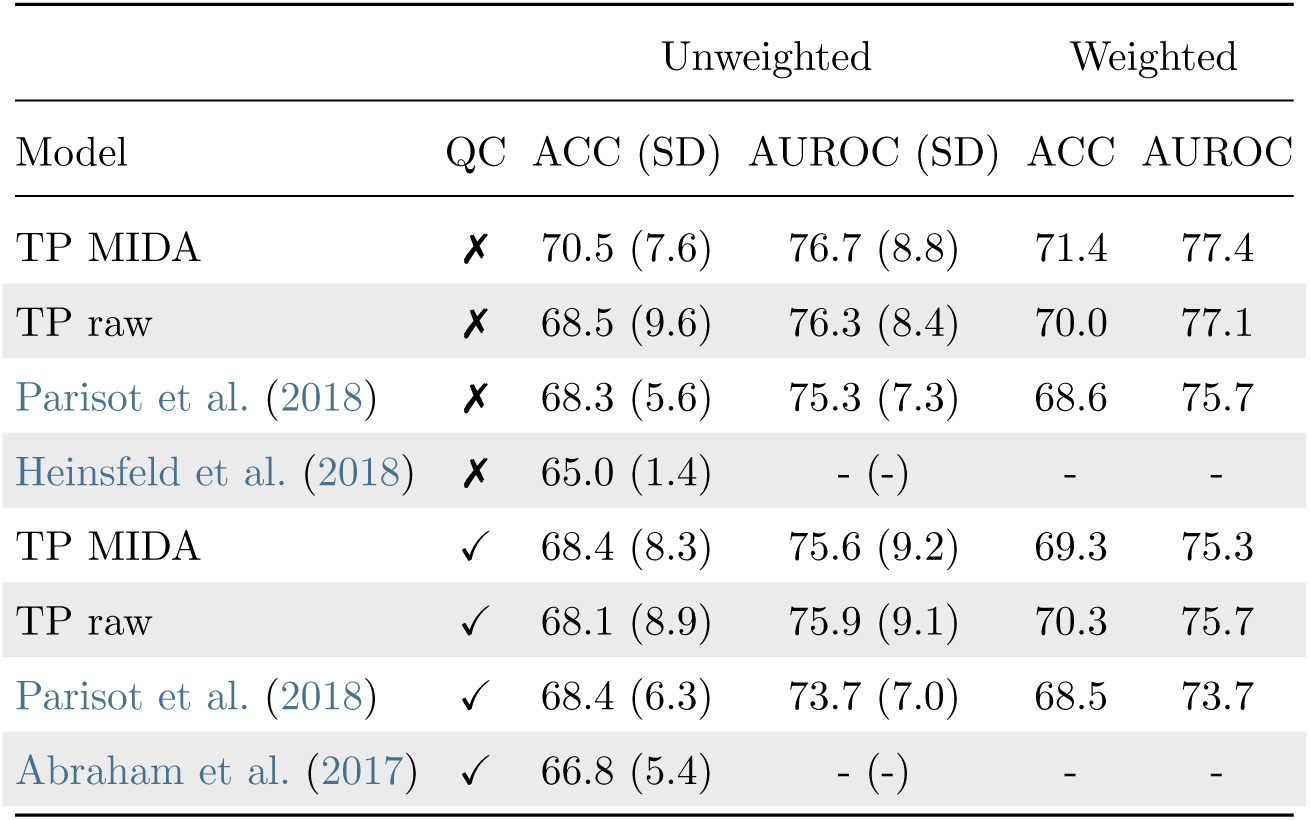
Inter-site comparison, i.e. leave one site out CV (LOSOCV). The notations in this table are the same as in Table 5.

| Model | Unweighted |  |  | Weighted |  |
| --- | --- | --- | --- | --- | --- |
|  | QC | ACC (SD) | AUROC (SD) | ACC | AUROC |
| TP MIDA | ✗ | 70.5 (7.6) | 76.7 (8.8) | 71.4 | 77.4 |
| TP raw | ✗ | 68.5 (9.6) | 76.3 (8.4) | 70.0 | 77.1 |
| Parisot et al. (2018) | ✗ | 68.3 (5.6) | 75.3 (7.3) | 68.6 | 75.7 |
| Heinsfeld et al. (2018) | ✗ | 65.0 (1.4) | - (-) | - | - |
| TP MIDA | ✓ | 68.4 (8.3) | 75.6 (9.2) | 69.3 | 75.3 |
| TP raw | ✓ | 68.1 (8.9) | 75.9 (9.1) | 70.3 | 75.7 |
| Parisot et al. (2018) | ✓ | 68.4 (6.3) | 73.7 (7.0) | 68.5 | 73.7 |
| Abraham et al. (2017) | ✓ | 66.8 (5.4) | - (-) | - | - |

In the 10-fold CV, our TP raw ridge (with phenotypic features, without domain adaptation) on a sample size of 1035 subjects achieves 71.6% in accuracy and 78.2% in AUROC, outperforming neural network based models (Parisot et al., 2018; Heinsfeld et al., 2018) by at least 1.2% in accuracy and 3.2% in AUROC. It obtained statistically significant increases in accuracy (3.4%, *p* = 0.02) and AUROC (3%, *p* = 0.05) at the 10% level w.r.t. to (Parisot et al., 2018) on 1035 subjects. In the 5×10-fold CV, we obtained *p*-values less than 1% when comparing the accuracy/AUROC of TP raw ridge on 1035 subjects with both 871/1035-subject variants of (Parisot et al., 2018). On 871 subjects, our TP raw LR achieves 70.1% in accuracy and 76.5% in AUROC, ourperforming (Parisot et al., 2018) and even TP MIDA LR, showing the effectiveness of the proposed tangent Pearson FC measure.

#### Inter-site comparison

Table 6 shows the inter-site performance comparison. We firstly observe that the weighted site accuracy and AUROC scores are generally higher than the unweighted results. This is expected because from the right panel of Fig. 5, sites with lower accuracy tends to have a smaller sample size. Secondly, our TP MIDA ridge (with phenotypic features) on 1035 samples achieves the highest (weighted) accuracy of 71.4%, improving the model by Parisot et al. (2018) on 871 and 1035 subjects by 2.9% and 2.8% in accuracy, respectively. Without domain adaptation, our TP raw ridge (with phenotypic features) also achieves a higher (weighted) accuracy (and AUROC) than (Parisot et al., 2018) when the sample size is 1035, an increase in accuracy by 1.5% and 1.4% w.r.t. (Parisot et al., 2018) on 871 and 1035 subject, respectively.

Across both sample sizes (871/1035), our proposed domain adaption and baseline models improve upon the models in Abraham et al. (2017) and Heinsfeld et al. (2018) in (unweighted) accuracy. On 1035 subjects, our TP MIDA ridge model has the highest accuracy of 70.5%, improving upon (Heinsfeld et al., 2018) by 5.5% and (Abraham et al., 2017) by 3.7%, and our baseline model TP raw ridge, improves over (Heinsfeld et al., 2018) and (Abraham et al., 2017) by 3.5% and 1.7%, respectively.

### 3.6. Extracting biomarkers

For the proposed FC measure, tangent Pearson, we study the respective ROI-ROI connections that have the most significant influence on the classification performance. These influential ROI-ROI connectivities can act as neurological biomarkers for researchers to investigate and further understand the difference in brain connectivity between ASD and TC.

To extract these biomarkers, we used the CC200 atlas to firstly define ROIs and the weights from the TP raw LR model (without phenotypic features) to indicate which ROI-ROI connections are most important for ASD/TC classification. The logistic regression classifier was used because it achieved the highest accuracy and AUROC for the tangent Pearson FC in the intra-site setting without phenotypic features. Phenotypic features were omitted because only ROI-ROI connections were of interest. The top five most positive and negative weights, for ASD and TC respectively, were extracted as described in Sec. 2.8.

The CC200 atlas is derived from the clustering of individual voxel BOLD time courses so the resulting atlas has no well-defined labels for the ROIs. To generate labels, we used the centre of mass for each ROI to locate the closest matching ROIs from other labelled brain atlases. In particular, we used the Harvard-Oxford brain atlas as a point of reference. Where no label could be found for a given CC200 ROI, we set its label to “None”.

Figure 6 shows the top 10 most important ROI-ROI connections for classifying subjects as ASD or TC, for illustration and further analysis in future studies. Red and blue connections give the top five most important ROI-ROI connections for classifying subjects as ASD and TC, respectively. The importance of each displayed connection is indicated by the hue of blue or red. Note that we omitted 180 of 200 ROIs from the figure to clearly show these connections.

**Figure 6:**
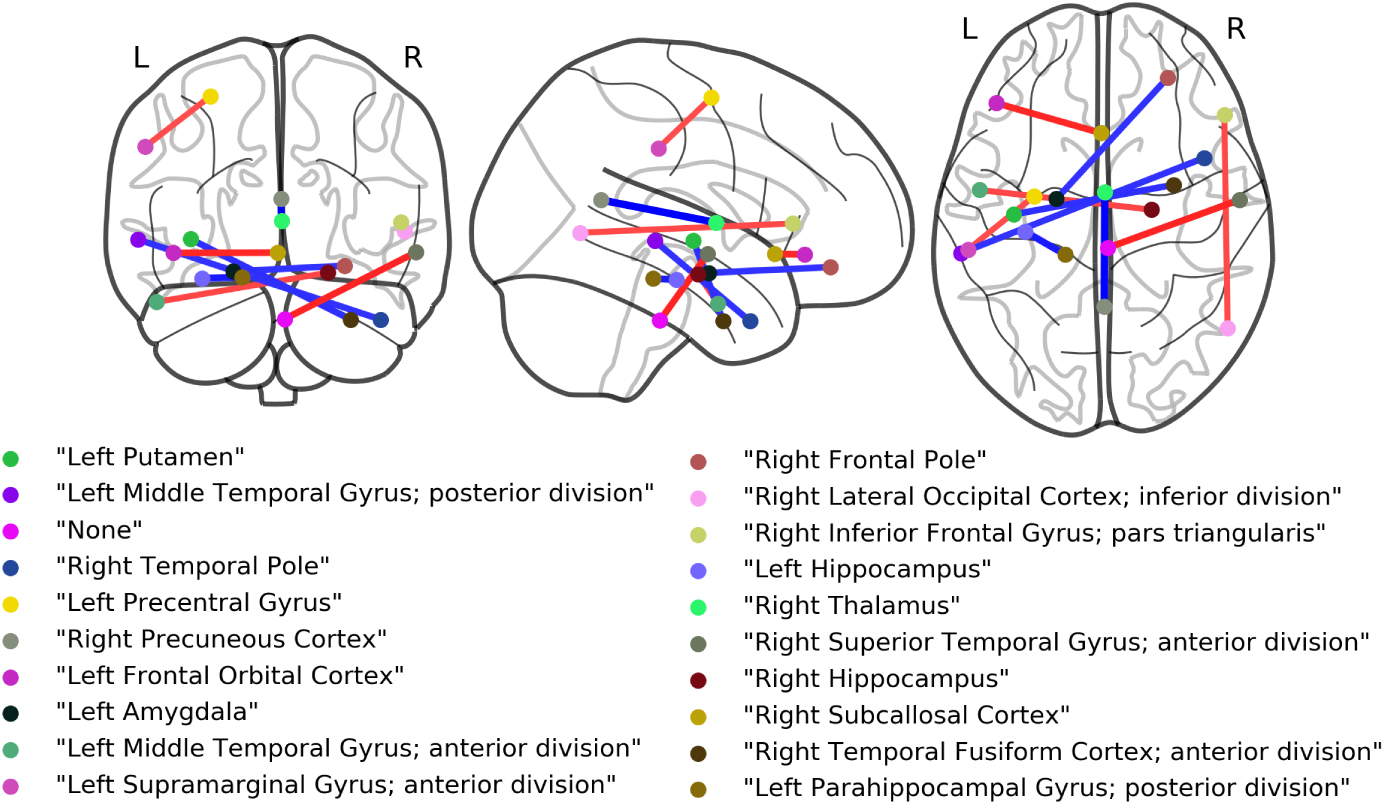
Biomarker visualisation. Extracted biomarkers using Python package Nilearn (Abraham et al., 2014). From left to right: the frontal, axial and lateral views of the brain are visualised. ‘L’ and ‘R’ correspond to the left and right hemisphere respectively.

## 4. Discussion

In this work, we proposed a second-order FC measure, tangent Pearson embedding, and investigated the minimisation of the dependence between ABIDE acquisition sites and FC features for improving autism classification. To firstly establish the significance of this study, we assessed the statistical dependence between acquisition sites and FC features, and then proceeded to assess the impact of removing site effects on the autism classification performance in the intra-site and inter-site experimental settings.

In comparison to a related study by Moradi et al. (2017) that targeted a problem of predicting autism severity scores with site-invariant cortical thickness data from four ABIDE sites, we targeted all possible 20 ABIDE sites with a sample of 1035 subjects. To assess the statistical dependence between subject FC features and acquisition sites, we used a kernel-based measure of statistical independence based on the empirical HSIC, which we found to be statistically insignificant at the 5% level with respect to a selection of three FC measures. In Figure 2, we further visualised the proposed second-order FC features with respect to acquisition sites, and the resulting plots revealed noticeable site specific representations of FC features, providing additional evidence of the site specificity of the rs-fMRI data in the ABIDE dataset highlighted in previous studies (Moradi et al., 2017; Parisot et al., 2018; Nielsen et al., 2013). This site specificity originates from variation in scanner type and experimental settings across sites, and has been hypothesised to be one of the limiting factors in training performant ASD classifiers on the ABIDE dataset (Parisot et al., 2018; Nielsen et al., 2013).

### Evaluation

For intra-site setting, *k*-fold CV is widely used in many studies. However, different random partitions can lead to significant variation of results, as seen by comparing the 10-fold CV and 5×10-fold CV results. Here we recommend *n* × *k*-fold CV, where the effect of random split settings is reduced, and the model evaluation is more stable. For inter-site setting, due to the differences in samples size across different sites, we recommend the weighted average accuracy, which represents 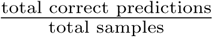 but unweighted average accuracy does not. In the rest of this section, our discussion will be based on the 5×10-CV and weighted leave-one-site-out CV results for intra-site and inter-site settings, respectively.

To study the impact of minimising unwanted dependence between acquisition site and subject FC features, we applied a domain adaptation approach, MIDA (Yan et al., 2017), in both intra-site and inter-site settings. In most cases, removing the dependence between site and FC features led to an improved performance in both settings (see Figures 3 and 4). MIDA-based models using the proposed tangent Pearson FC outperformed recent SOTA approaches (Parisot et al., 2018; Heinsfeld et al., 2018), achieving new SOTA performance of 72.7% in intra-site accuracy (AUROC: 77.9%), and 71.4% in inter-site accuracy (AU-ROC: 77.4%), corresponding to increases of 4.8% and 2.9% (AUROC: 4.6% and 3.7%) w.r.t. (Parisot et al., 2018), respectively. The results reported in this paper highlight the value of minimising the dependence between FC features and acquisition sites for improving autism classification. By removing acquisition site effects in FC features via a site-invariant subspace projection, classifiers trained on site-invariant features can extract more discriminative features for improving autism classification.

However, there are two limitations that are apparent when using MIDA to minimise site dependence. 1) Firstly, in a few cases, particularly with the tangent FC, when MIDA is applied, there is a degradation in intra/inter-site accuracy and AUROC scores relative to baseline models. A potential cause for such performance degradation is the difficulty in preserving relationships between projected subject FC features and target labels (ASD or TC). Though the variance in the original FC features can be preserved with MIDA, the alignment between projected features and target labels may not be (fully) preserved. This is especially important for FC features derived from rs-fMRI data for autism classification since the underlying signal defining autism is not well marked, due to the heterogeneity of ASD. Therefore, applying a technique that uses the training data labels to align subject FC features without overfitting may help unlock more potential from the ABIDE dataset in multi-site autism classification. 2) Secondly, since MIDA is a transductive learning method, adding new subjects to the experimental dataset would require a new domain invariant subspace to be computed that accounts for the new data before predictions can be made. Another research focus is therefore applying or developing an inductive domain adaptation approach that alleviate this problem.

### ‘Low-quality’ samples in domain adaptation

For both intra-site and intersite settings, we observed accuracy improvement obtained by domain adaptation (MIDA) when including more ‘low-quality’ samples in training. In contrast to such positive effect for MIDA-based models, including them has much less effect on the classification accuracy of other models. On one hand, these samples may be helpful in estimating the site data distribution so that MIDA can extract better domain-invariant features. On the other hand, their phenotypic features are not necessarily low quality and can also contribute to the autism prediction. The proposed tangent Pearson embedding FC measure has a simple implementation built on existing FC measures, Pearson correlation and tangent embedding. It has been shown to be a powerful alternative. Without the application of MIDA, we observed that this new second-order FC measure can outperform previous SOTA methods when supplemented with phenotypic features and a linear classifier. In particular, we achieved the highest accuracy score of 70.9% (AUROC: 78.0%) in the intra-site experiment, and an accuracy of 70.0% (AUROC: 77.1%) in the inter-site setting, improving upon (Parisot et al., 2018) by 3.0% and 1.6% in accuracy (4.7% and 3.4% in AUROC), respectively. This makes our proposed FC measure particularly attractive, considering the results from those SOTA methods employing complex neural networks that take a long time to train (e.g. 32 hours for (Heinsfeld et al., 2018)). The linear classifiers investigated in this study are capable of achieving improved results with only several minutes of training when leveraging the proposed second-order FC features and phenotypic information.

Lastly, the proposed second-order FC measure and adopted MIDA approach affect only the feature representation of each subject to improve multi-site autism classification. The obtained performance is mostly similar across three different classifiers (see Figures 3 and 4). Therefore, we hypothesise that to improve multi-site autism classification, it is important to 1) design more powerful FC measures or other measures from raw fMRI data, and 2) directly target and remove site agglomerative effects.

## 5. Conclusions

This paper aims to improve multi-site autism classification. We proposed a new second-order functional connectivity (FC) measure called tangent Pearson embedding for more discriminative FC features, and applied a site-dependence-minimisation domain adaptation approach to tackle the heterogeneity in the multi-site ABIDE database. We confirmed the significance of this study via a statistical independence assessment between acquisition sites and FC features. The intra- and inter-site classification results show that models with the proposed FC measure and site-dependence minimisation in combination with pheonotypic features have outperformed the state-of-the-art methods, with detailed ablation studies and biomarker visualisation.

## Acknowledgement

This work was partially supported by the UK Engineering and Physical Sciences Research Council [EP/R014507/1], and the National Natural Science Foundation of China [81671772]. The authors would like to thank all the sites and investigators who have shared data through ABIDE.

## Footnotes

1 http://preprocessed-connectomes-project.org/abide/

2 Available at https://github.com/parisots/population-gcn

